# A malaria transmission model with seasonal mosquito life-history traits

**DOI:** 10.1101/377184

**Authors:** Ramsès Djidjou-Demasse, Gbenga J. Abiodun, Abiodun M. Adeola, Joel O. Botai

## Abstract

In this paper we develop and analyse a malaria model with seasonality of mosquito life-history traits: periodic-mosquitoes per capita birth rate, -mosquitoes death rate, -probability of mosquito to human disease transmission, -probability of human to mosquito disease transmission and -mosquitoes biting rate. All these parameters are assumed to be time dependent leading to a nonautonomous differential equation systems. We provide a global analysis of the model depending on two thresholds parameters 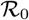 and 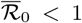 (with 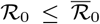). When 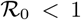, then the disease-free stationary state is locally asymptotically stable. In the presence of the human disease-induced mortality, the global stability of the disease-free stationary state is guarantied when 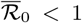. On the contrary, if 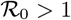, the disease persists in the host population in the long term and the model admits at least one positive periodic solution. Moreover, by a numerical simulation, we show that a subcritical (backward) bifurcation is possible at 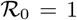. Finally, the simulation results are in accordance with the seasonal variation of the reported cases of a malaria-epidemic region in Mpumalanga province in South Africa.

## 1 Introduction

Malaria is a deadly disease caused by infection with *Plasmodium* protozoa transmitted by an infective female *Anopheles* mosquito. Globally, an estimated 216 million malaria cases was recorded in 2016 which is an increase of about 5 million cases over 2015 [47]. In the same year, almost 445,000 individuals lost their lives to the life-threatening disease [47].

Despite more than a century of research, there is a dearth of information on the mechanistic link between environmental variables, such as temperature and malaria risk [29, 38, 7]. Temperature is fundamentally linked to malaria mosquito and parasite vital rates, and understanding the role of temperature in malaria transmission is particularly important in light of climate change. Using mathematical model, several attempts have been made to highlight some of the importance of climate variables on malaria transmission. [33] built nonlinear thermal-response models to understanding the effects of current and future temperature regimes on malaria transmission. The models, which include empirically derived nonlinear thermal responses, predicts optimal malaria transmission at 25°C (6°C lower than previous models).

[21] proposed an ordinary differential equation (ODE) compartmental model for the spread of malaria with susceptible-infectious-recovered (SIRS) pattern for humans and a susceptible-infectious (SI) pattern for mosquitoes with mosquitoes periodic birth rate and death rate. More recently, [3] developed and analysed a comprehensive mosquito-human dynamical model. The model was validated by [2] over KwaZulu-Natal province – one of the epidemic provinces in South Africa. Several other studies [20, 30, 39, 4, 37, 1, 22, 12, 26] have explored the impacts of environmental variables on malaria transmission and mosquito abundance. We reference the work of [43] with periodic mosquito per capita death and birth rate; recent works of [11] with periodic mosquito biting rate and [36] for a model assessing the impact of the differences in indoor vs outdoor environments on the efficacy of malaria control. [37] assess the impact of variability in temperature and rainfall on the transmission dynamics of malaria over KwaZulu-Natal, South Africa and found that incorporating host age-structure and reduced susceptibility due to prior malaria infection has marginal effect on the transmission dynamics of the disease. Similarly, [6] presented a new mechanistic deterministic model to investigate the impact of temperature variability on malaria transmission over East, West, South and Central Africa. With temperature between 16–34°C, their analysis identified mosquito carrying capacity, transmission probability per contact for susceptible mosquitoes and human recruitment rate as the most sensitive parameters. However, many other mosquito life-history traits (including larval development rate, larval survival, adult survival, biting rate, fecundity, and vector competence) are well known to have seasonal variation [33, 22]. This present study aims to (i) proposed and analyze a human-mosquito malaria transmission model including all these life-history traits with periodic variation and (ii) validate the model proposed over a malaria-epidemic regions in Mpumalanga province in South Africa.

The model proposed in this paper divides the human population into four classes: susceptible-exposed-infectious-recovered (SEIRS) and mosquitoes population into three classes: susceptible-exposed-infectious (SEI). The SEIRS pattern for humans and SEI pattern for mosquito model have been also proposed by Chitnis and collaborators [18, 19]. Human migration is present throughout the world and plays a large role in the epidemiology of diseases, including malaria. In many parts of the developing world, there is rapid urbanization as many people leave rural areas and migrate to cities in search of employment. We include this movement as a constant immigration rate into the human susceptible class. We make a simplifying assumption that there is no immigration of recovered humans and also include the direct infectious-to-susceptible recovery as in the model of [35].

This work is organized as follows: In Section 2, we fully describe the malaria seasonal model studied in this paper as Section 3 describes the main results. The discussion and numerical simulations illustrating main results are given in Sections 4. Section 5 is devoted for deriving preliminary results and remarks that will be used to study the long-term behavior of the problem. Section 6 is concerned with the proof of the main results that, roughly speaking, state that when some thresholds (explicitly expressed using the parameters of the system) 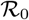 and 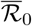 (with 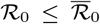) are such that: (i) when 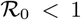 the disease-free stationary state is locally (not necessarily globally) asymptotically stable; (ii) when 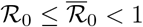 the disease-free stationary state is then globally asymptotically stable and the disease certainly die out from the host population; and (iii) when 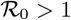 the disease persists in the host population in the long term and the model admits at least one positive periodic solution.

## 2 The malaria seasonal model

The model sub-divides the total human population at time *t*, denoted by *N*_*h*_(*t*), into the following sub-populations of susceptible *S*_*h*_(*t*), exposed (infected but not infectious) *E*_*h*_(*t*), infectious *I*_*h*_(*t*) and recovered individuals with temporary immunity *R*_*h*_(*t*). So that *N*_*h*_(*t*) = *S*_*h*_(*t*) + *E*_*h*_(*t*) + *I*_*h*_(*t*) + *R*_*h*_(*t*).

The total vector (mosquito) population at time *t*, denoted by *N*_*v*_(*t*), is sub-divided into susceptible *S*_*v*_(*t*), exposed *E*_*v*_(*t*) and infectious mosquitoes *I*_*v*_ (*t*). Thus, *N*_*v*_(*t*) = *S*_*v*_(*t*) + *E*_*v*_(*t*) + *I*_*v*_(*t*).

Susceptibles individuals are recruited at a constant rate Λ_*h*_. We define the force of infection from mosquitoes to humans by (*αβ*_1_(*t*)*θ*(*t*)*I*_*v*_/*N*_*h*_) as the contact probability between human and mosquito *α*, the probability that the mosquito is infectious *I*_*v*_/*N*_*v*_, the number of mosquito bites per human per time *θ*(*t*)*N*_*v*_/*N*_*h*_ and the probability of disease transmission from the mosquito to the human *β*_1_(*t*). Then infected individuals move to the exposed class at a rate (*αβ*_1_(*t*)*θ*(*t*)*I*_*v*_*S*_*h*_/*N*_*h*_). The natural death rate of human is *μ*_*h*_. The rate of progression from exposed class to infectious individuals class is *σ*_*h*_ while infectious individuals recovered due to treatment at a rate *γ*_*h*_. The infectious humans after recovery without immunity become immediately susceptible again at rate (1 − *r*), where *r* is the proportion of infectious humans who recovered with temporary immunity. Recovered individual loses immunity at a rate *k*_*h*_.

Susceptible mosquitoes are generated at a per capita rate *b*_*v*_(*t*) at time *t* and acquire malaria through contacts with infectious humans with the force of infection (*β*_2_(*t*)*αθ*(*t*)*I*_*h*_/*N*_*h*_) as the product of the probability of disease transmission from human to the mosquito *β*_2_(*t*), the human-mosquito contact probability *α* and the probability that human is infectious *I*_*h*_/*N*_*h*_. Hence, newly infected mosquitoes are moved into the exposed class at a rate (*β*_2_(*t*)*αθ*(*t*)*I*_*h*_*S*_*v*_/*N*_*h*_) and progress to the class of infectious mosquitoes at a rate *σ*_*v*_. Mosquitoes are assumed to suffer death at rate (*μ*_*v*_(*t*) + *κ*_*v*_(*t*)*N*_*v*_), either due to natural causes at rate *μ*_*v*_(*t*) or to the density-dependent death at rate *κ*_*v*_(*t*)*N*_*v*_ at time *t*.

The non-autonomous model has time-dependent *ω*-periodic coefficients which account for the environmental variations in the infectivity of both human and mosquitoes populations, the birth rate of the mosquitoes population, the biting rate of the mosquitoes population and the death rates of mosquitoes. Setting 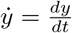, the resulting system of equation is shown below:

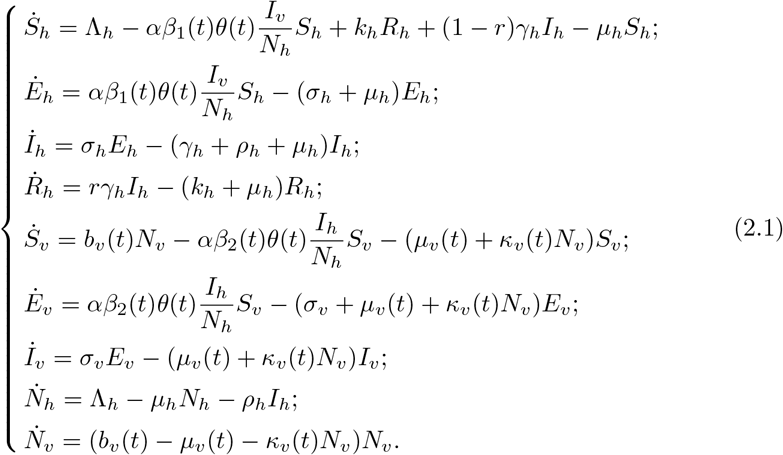

For convenience, we work with fractional quantity instead of actual populations by scaling the population of each by the total species population. We let

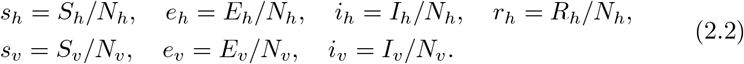

Differentiating the scaling equations (2.2) and solving for the derivatives of scaled variables, we obtain for example

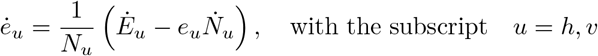

and so on for the other variables.

This creates a new seven-dimensional system of equations with one dimension for the total population variable, *N*_*h*_, and five dimensions for the fractional population variables, *e*_*h*_, *i*_*h*_, *r*_*h*_, *e*_*v*_, and *i*_*v*_. For convenience we still use *s*_*h*_ = *S*_*h*_, *e*_*h*_ = *E*_*h*_, *i*_*h*_ = *I*_*h*_, *s*_*v*_= *S*_*v*_, *e*_*v*_ = *E*_*v*_ and *i*_*v*_ = *I*_*v*_. We then have

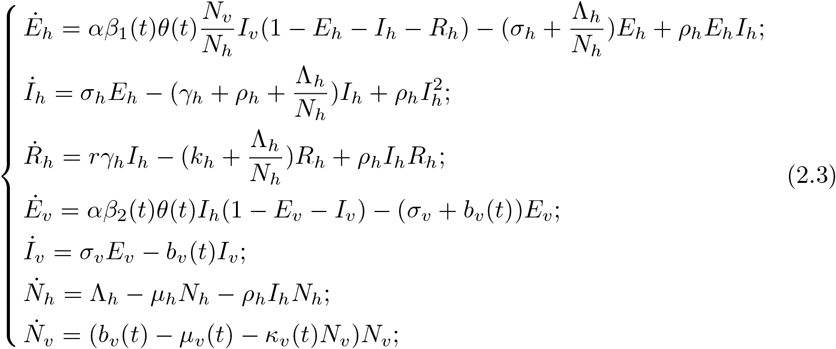

and

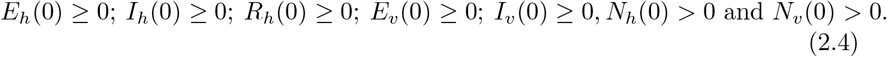

Variables and parameters of the model are resumed in Table 1. In what follow we shall discuss the asymptotic behavior of system (2.3)-(2.4) and we will make use the following assumptions.

**Table 1:** Parameter description

| Constant parameters |  |
| --- | --- |
| $\Lambda_h$ | human recruitment rate. $Human \times Time^{-1}$. |
| $\mu_h$ | human per capita death rate. $Time^{-1}$. |
| $\alpha$ | human-mosquito contact probability. *Dimensionless*. |
| $k_h$ | rate of loss of immunity. $Time^{-1}$. |
| $\gamma_h$ | human recovery rate. $Time^{-1}$. |
| $r$ | Probability of recovered with temporary immunity. *Dimensionless*. |
| $\sigma_h$ | progression rate from exposed class to infectious individuals. $Time^{-1}$. |
| $\sigma_v$ | progression rate from exposed class to infectious mosquitoes. $Time^{-1}$. |
| $\rho_h$ | human disease induce mortality rate. $Time^{-1}$. |
| ω-Periodic parameters | |
| $b_v(t)$ | mosquitoes per capita birth rate. $Time^{-1}$. |
| $\mu_v(t)$ | mosquitoes per capita death rate. $Time^{-1}$. |
| $\kappa_v(t)$ | Density-dependent part of the death rate for mosquitoes. $Mosquito^{-1} \times Time^{-1}$ |
| $\beta_1(t)$ | probability of mosquito to human disease transmission. *Dimensionless*. |
| $\beta_2(t)$ | probability of human to mosquito disease transmission. *Dimensionless*. |
| $\theta(t)$ | mosquitoes biting rate. $Human \times Mosquito^{-1} \times Time^{-1}$ |

### Assumption 2.1

1. *We assume that*, Λ_*h*_, *μ*_*h*_, *α*, *k*_*h*_, *γ*_*h*_, *r*, *σ*_*h*_ and *σ*_*v*_ *are positives constants with the exception of the disease-induced death rate ρ*_*h*_, *which is non-negative constant. The functions b*_*v*_(.), *μ*_*v*_(.), *κ*_*v*_(.), *β*_1_(.), *β*_2_(.) *and θ*(.) *are ω-periodic and belong to* 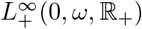.
2. *max*_*t*∈[0,*ω*]_ (*b*_*v*_(*t*) − *μ*_*v*_(*t*)) > 0 *and* min_*t*∈[0,*ω*]_ *κ*_*v*_(*t*) > 0.

Biologically, item 1. of Assumption 2.1 is a quite natural assumption on demographical, epidemiological and biological model parameters and item 2. means that the maximum of mosquito’s natural growth rate is positive. For notational convenience, we define the average of any *ω*-periodic function *z* by setting 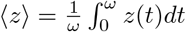.

## 3 Main results

In what follows, we characterize the disease-free periodic state and introduce the basic reproduction number 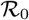 for system (2.3) according to general procedure presented in [45, 31] and references therein.

### 3.1 The disease-free periodic state

In the absence of disease, total populations of human and vectors *N*_*h*_ and *N*_*v*_ are such that

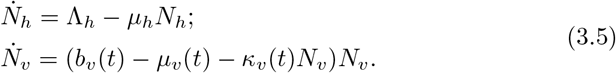

By Assumption 2.1, the disease-free *ω*-periodic state of system (2.3) is 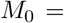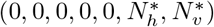; where 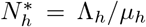 and 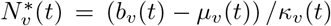. Moreover, 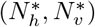 is the unique positive *ω*-periodic solution of System (3.5), which is globally attractive in 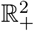 (see section 5.1 for details of this proof).

### 3.2 The basic reproduction number 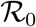

In the sequel, let us introduce the *ω*-periodic function 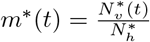, the ratio of mosquitoes to human population. Then, the equation for exposed and infectious for both human and mosquitoes populations of the linearized system of model (2.3) at *M*_0_ writes

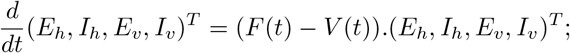

where

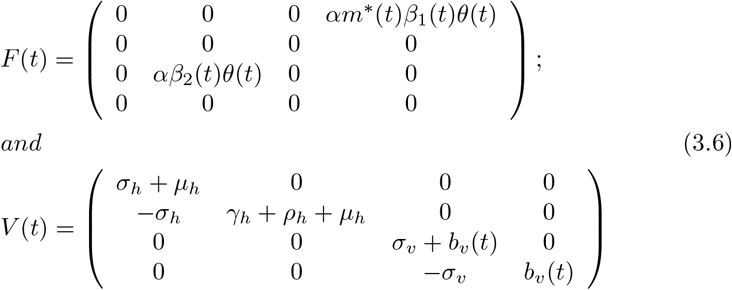

Let Φ_*V*_(*t*) and *ρ*(Φ_*V*_(*ω*)) be the monodromy matrix of the linear *ω*-periodic system 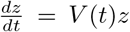 and the spectral radius of Φ_*V*_(*ω*), respectively. Assume *Y*_*V*_(*t*, *s*), *t* ≥ *s*, is the evolution operator of the linear *ω*-periodic system

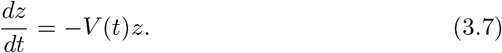

That is, for each *s* ∈ ℝ, the 4 × 4 matrix *Y*(*t*, *s*) satisfies

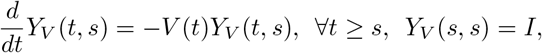

where *I* is the 4 × 4 identity matrix. Thus, the monodromy matrix Φ_−*V*_(*t*) of (3.7) is equal to *Y*_*V*_(*t*, 0) for *t* ≥ 0.

Now, we deal with disease-free equilibrium invasion process [44] and references therein. Let *ϕ*(*s*) the initial distribution of infectious individuals. Then *F*(*s*)*ϕ*(*s*) is the rate of new infections produced by the infected individuals who were introduced at time *s*. Given *t* ≥ *s*, then *Y*_*V*_ (*t*, *s*)*F*(*s*)*ϕ*(*s*) gives the distribution of those infected individuals who were newly infected at time *s* and remain in the infected compartments at time *t*. It follows that

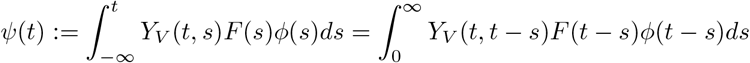

is the distribution of accumulative new infections at time *t* produced by all those infected individuals *ϕ*(*s*) introduced at time previous to *t*.

Let 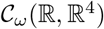 be the ordered Banach space of all *ω*-periodic functions from ℝ to ℝ^4^ which is equipped with the maximum norm ∥.∥ and the positive cone 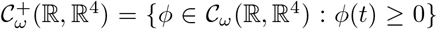. Then we can define a linear operator 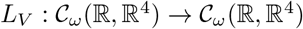 by

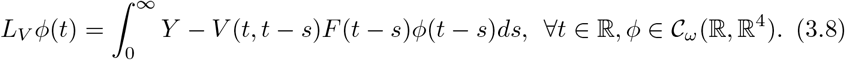

Following [45], we call *L* the next generation operator, and define the basic reproduction number as 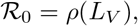, the spectral radius of *L*_*V*_. In the context of this work, the computation of 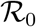 of model (2.3) is based on Lemma 5.3, item (ii). Other approximate formula and numerical methods can be founded in [10] and references cited therein.

In the special case of *β*_1_(*t*) ≡ *β*_1_, *β*_2_(*t*) ≡ *β*_2_, *b*_*v*_(*t*) ≡ *b*_*v*_, *μ*_*v*_(*t*) ≡ *μ*_*v*_, *κ*_*v*_(*t*) ≡ *κ*_*v*_ and *θ*(*t*) ≡ *θ* ∀*t* ≥ 0, we obtain *F*(*t*) ≡ *F* and *V* (*t*) ≡ *V*. By [44], the basic reproduction number is:

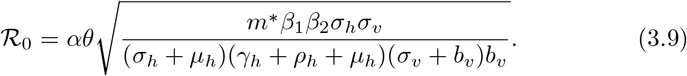

As pointed in [18], The original definition of the reproductive number of the Ross-Macdonald model [8] and the Ngwa and Shu model [35], is equivalent to the square of this 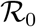. Anderson and Ngaw use the traditional definition of the reproductive number, which approximates the number of secondary infections in humans caused by one infected human, while the 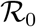 used here is consistent with the definition given by the next generation operator approach [44] which approximates the number of secondary infections due to one infected individual (be it human or mosquito). Moreover, the number of new infections in humans that one human causes through his/her infectious period is 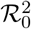, not 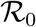. Because this defination of 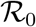 (3.9) is based on the next generation operator approach, it counts the number of new infections from one generation to the next. That is, the number of new infections in mosquitoes counts as one generation. We further define the average basic reproduction number (according to the periodic parameters)

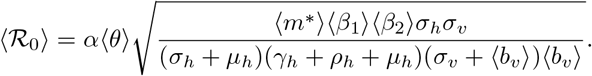

In general, 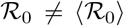. For example, in a tuberculosis model it was shown, in [31], that 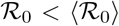, and in Dengue fever model it was shown in [45] that 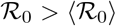. We can also consult [23] for more details.

### 3.3 The threshold dynamics

To deal with model (2.3), some notations will be given. Let us identify *x*_*h*_ together with (*E*_*h*_, *I*_*h*_, *R*_*h*_, *N*_*h*_)^*T*^ and *x*_*v*_ together with (*E*_*v*_, *I*_*v*_, *N*_*v*_)^*T*^ and set 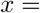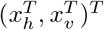. Define

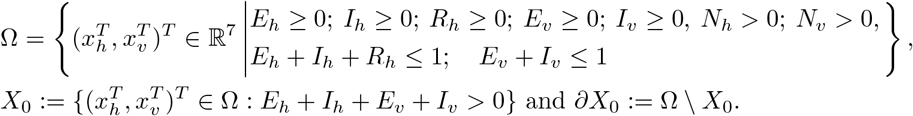

Using the above notations the main result of this work is the following theorem.

#### Theorem 3.1

*Let Assumption 2.1 be satisfied.*

i. *The disease-free equilibrium M*_0_ *for System* (2.3)-(2.4) *is locally asymptotically stable if* 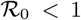 *and unstable if* 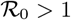.
ii. *If* 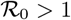, *then system* (2.3)-(2.4) *is uniformly persistence with respect to the pair* (*X*_0_, *∂X*_0_), in the sense that there exists *δ* > 0, such that for any *x*_0_ ∈ *X*_0_ *we have*,

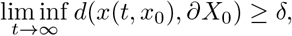

*and system* (2.3) *admits at least one positive periodic solution; where x*(*t*, *x*_0_) *is the unique solution of* (2.3) *with x*(0, *x*_0_) = *x*_0_.

Details of this proof are given in section 6. Theorem 3.1 states that there are at least two possible realistic equilibrium: one where the disease die out from the host population and the other, if 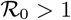, where the disease persists (with periodical behaviour) in the host population. So, 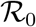 is a threshold parameter which *partially* determines the epidemic behaviour of Model (2.3). Indeed, generally in epidemiological models, bifurcations at the threshold 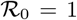 tend to be supercritical (or forward), meaning that a positive endemic equilibria exist for 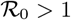 near the bifurcation point. In model (2.3) we prove that the bifurcation is supercritical in the absence of disease-induced death (*ρ*_*h*_ = 0). In general case (*ρ*_*h*_ > 0), the local stability result of the disease-free equilibrium provided by Theorem 3.1 means that, in the context of model (2.3), there can be stable endemic equilibrium when 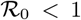 (see [18] for more discussions on backward bifurcation in some epidemiological model).

However, the global stability of the disease-free equilibrium can be derive when *ρ*_*h*_ > 0 with some additional hypothesis. For that ends, let *ρ*_*h*_ > 0 be small enough. We introduce the following matrices

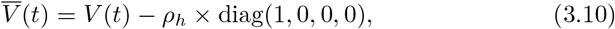

wherein diag(*u*) is set for a diagonal matrix which diagonal elements are given by *u*. Let 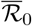 be the spectral radius of the operator 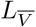 defined by (3.8) and wherein the matrix *V* is replaced by 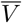. We then have the following result of the global stability of the disease-free equilibrium.

#### Theorem 3.2

*Let Assumption 2.1 be satisfied and let ρ*_*h*_ > 0 *be small enough such that ρ*_*h*_ < *μ*_*h*_ + *γ*_*h*_. *Then, the disease-free equilibrium M*_0_ *for System* (2.3)-(2.4) *is globally asymptotically stable if* 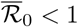.

Details on the proof of Theorem 3.2 are given in section 6.2. Moreover, in section 6.3 we show that in the absence of disease-induced death (*ρ*_*h*_ = 0), 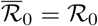 and 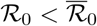 when *ρ*_*h*_ > 0.

## 4 Numerical results and discussion

In this section, we provide some numerical simulations to support and discuss our analytical conclusions. For that end, we start with some reasonable assumption on the density dependent death-rate of mosquitoes *κ*_*v*_(*t*). Due to the density dependent death rate *κ*_*v*_ (*t*), we assume that the vector population *N*_*v*_ has small seasonal fluctuations at equilibrium. Therefore, *κ*_*v*_(*t*) ≃ *C*_0_ (*b*_*v*_(*t*) − *μ*_*v*_(*t*)), wherein *C*_0_ is a positive constant. Doing that, the ratio 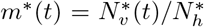 of mosquitoes to human population is quasi constant and we have 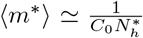, *i.e.* 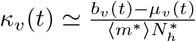.

The temperature-dependent parameters of model (2.1) are defined using thermal response functions describe in [33]. Our simulations are based on the daily climate temperature of Nkomazi from 1997 to 2005.

### Thermal-response curves

**A** collection of data to derive functions relating vector and parasite parameters to temperature was updated by [33]. As all rate parameters in the temperature-dependent are expected to be unimodal with respect to temperature, [33] fit quadratic and Briére functions to each life-history parameter, as well as a linear function for comparison (Table 2). The Briére function is a left-skewed unimodal curve with three parameters, which represent the minimum temperature, maximum temperature and a rate constant[16]. The unimodal functions are defined as Briére [*c*(*T*_0_ − *T*(*t*))(*T*_*m*_ − *T*(*t*))^1/2^] and quadratic [*qT*^2^(*t*) + *rT*(*t*) + *s*], where *T*(*t*) is temperature in degrees Celsius at time *t*. Constants *c*, *T*_0_, *T*_*m*_, *q*, *r* and *s* are fitting parameters. In this paper, mosquitoes per capita birth rate (*b*_*v*_), per capita death rate (*μ*_*v*_), infectivity (*β*_2_) and biting rate (*θ*) are estimated from thermal performance curves and summarized in Table 2.

**Table 2:** The relationships between temperature and the mosquito and parasite life-history traits that determine malaria risk. Thermal performance curves were fitted to the data assuming Briére [*c*(*T*_0_ − *T*(*t*))(*T*_*m*_ − *T*(*t*))^1/2^], **B**, or Quadratic [*qT*^2^(*t*) + *rT*(*t*) + *s*], **Q**, functions; in which *T*(*t*) is temperature (°*C*) at time t. Standard deviations for the parameters are listed in parentheses alongside parameter values. (see [33] and references therein).

| Parameters | Definition | Fit | Fit parameters (standard deviation) |  |  |  |
| --- | --- | --- | --- | --- | --- | --- |
| $\theta$ | Biting rate | **B** | $c = 0.000203(0.0000576)$ | $T_m = 42.3(3.53)$ | $T_0 = 11.7(2.47)$ | |
| $\beta_1 * \beta_2$ | Vector competence | **Q** | $q = -0.54(0.18)$ | $r = 25.2(9.04)$ | $s = -206(108)$ | |
| $e^{-\mu_v}$ | Daily adult survival probability | **Q** | $q = -0.000828(0.0000519)$ | $r = 0.0367(0.00239)$ | $s = 0.522(0.0235)$ | |
| Mdr | Mosquito development rate | **B** | $c = 0.000111(0.00000954)$ | $T_m = 34(0.000106)$ | $T_0 = 14.7(0.831)$ | |
| pEA | Egg-to-adult survival probability | **Q** | $q = -0.00924(0.00123)$ | $r = 0.453(0.0618)$ | $s = -4.77(0.746)$ | |
| EFD | Egg laid per adult female per day | **Q** | $q = -0.153(0.0307)$ | $r = 8.61(1.69)$ | $s = -97.7(22.6)$ | |
| PDR | Parasite development rate | **B** | $c = 0.000111(0.0000161)$ | $T_m = 34.4(0.000176)$ | $T_0 = 14.7(1.48)$ | |
| Using the above notations, the mosquitoes per capita birth rate $b_v$ is given by | | | | | | |
| $b_v(t) = EFD(t) * Mdr(t) * pEA(t) * e^{-\mu_v(t)}$. | | | | | | |

### Nkomazi (South Africa) climate data

For all simulations, we incorporate the daily climate data (temperature) of Nkomazi from 1997 to 2005 into our model to estimate time dependent parameters of model (2.1), namely *b*_*v*_, *μ*_*v*_, *β*_2_ and *θ* (Figure 1). The temperature data were extracted from the National Centers for Environmental Prediction (NCEP) Climate Forecast System Reanalysis (CFSR). The 6-hourly climate dataset was converted to daily with 0.5°×0.5° resolution for the purpose of this study. Conversely, the malaria data sourced from the provincial Integrated Malaria Information System (IMIS) of the malaria control program in the Mpumalanga Provincial Department of Health, was obtained from the South African Weather Service (SAWS) through its collaborative research with the University of Pretoria Institute for Sustainable Malaria Control (UP ISMC). The locally recorded cases with minimal imported cases were extracted from Nkomazi – a local municipality in Mpumalanga province (one of the epidemic provinces in South Africa). In the province, malaria distribution is mainly in Nkomazi, Bushbuckridge, Mbombela, Umjindi and Thaba Chewu local municipalities, with suitable climate conditions for malaria transmission [41, 5]. Of all the municipalities, Nkomazi has been identified as the most epidemic region in the province [41, 5]. The province recorded high malaria cases between 1998 and 2002 (Figure 5). Similar increase in cases were recorded across other epidemic provinces in South Africa during this period [41, 5]. Although studies show that most of the cases were imported [27]. For this reason, several malaria control measures were introduced to the provinces, which is traceable to the reduction in transmission after 2002 [41, 5].

**Figure 1:**
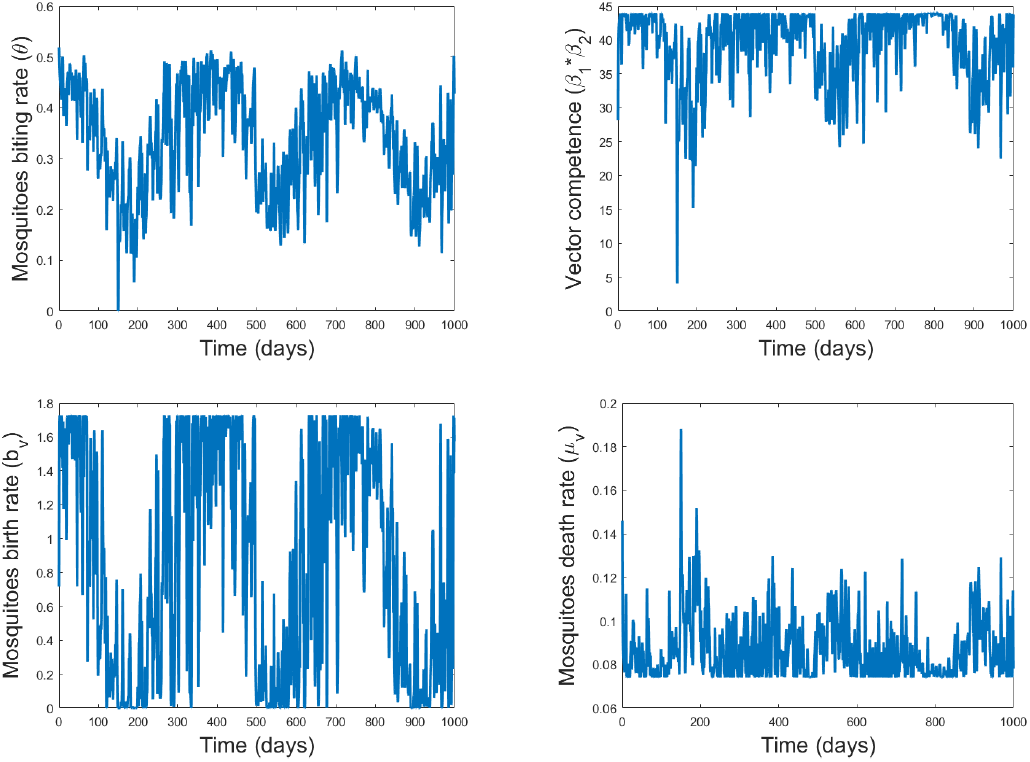
Time performance curves of mosquitoes traits using daily temperature of Nkomazi (South Africa) and thermal performance curves summarized in Table 2.

### The risk of outbreak in Nkomazi is underestimated when using the average basic reproduction number

Here we take *ρ*_*h*_ = 0.01, *γ*_*h*_ = 0.01, *r* = 0.9 and Λ_*h*_ = 0.05. We choose the total number of human and mosquitoes at initial time (*t* = 0) to be *N*_*h*_(0) = 1000 and *N*_*v*_ (0) = 2000 respectively. Two initial data of the model are considered and given in Table 3. Using the other parameters given by Tables 2 and 3, then by numerical computation, we get the curve of the basic reproduction number 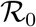 (applying Lemma 5.3 item (ii)) and the curve of the average basic reproduction number 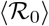 with respect to the mosquitoes contact rate *α* in Figure 2A. We can see that the average basic reproduction number 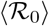 is always lower than the basic reproduction number 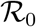 with *α* ranging from 0 to 0.5. Therefore, using 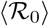 rather than 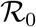 underestimates the outbreak of the disease in Nkomazi. Indeed, taking *α* = 0.2, numerical computation leads to 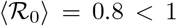, suggesting that there is no epidemic into the host population, and 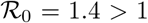, suggesting that there is an epidemic into the host population (Figure 2B-C). We have also computed the threshold parameter 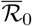 and we can note that 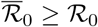.

**Table 3:**
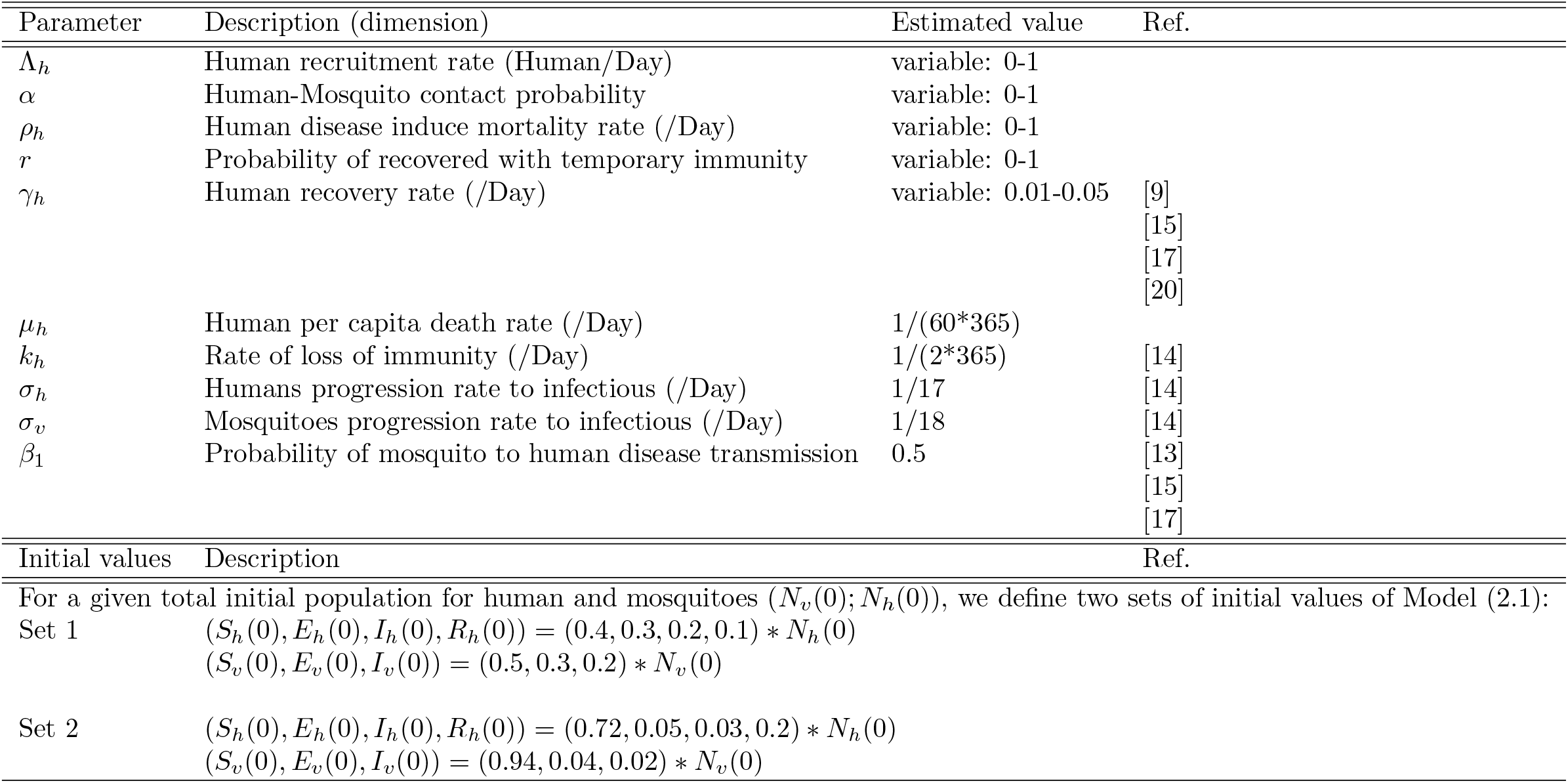
Initial data, parameters values and range of the model

**Figure 2:**
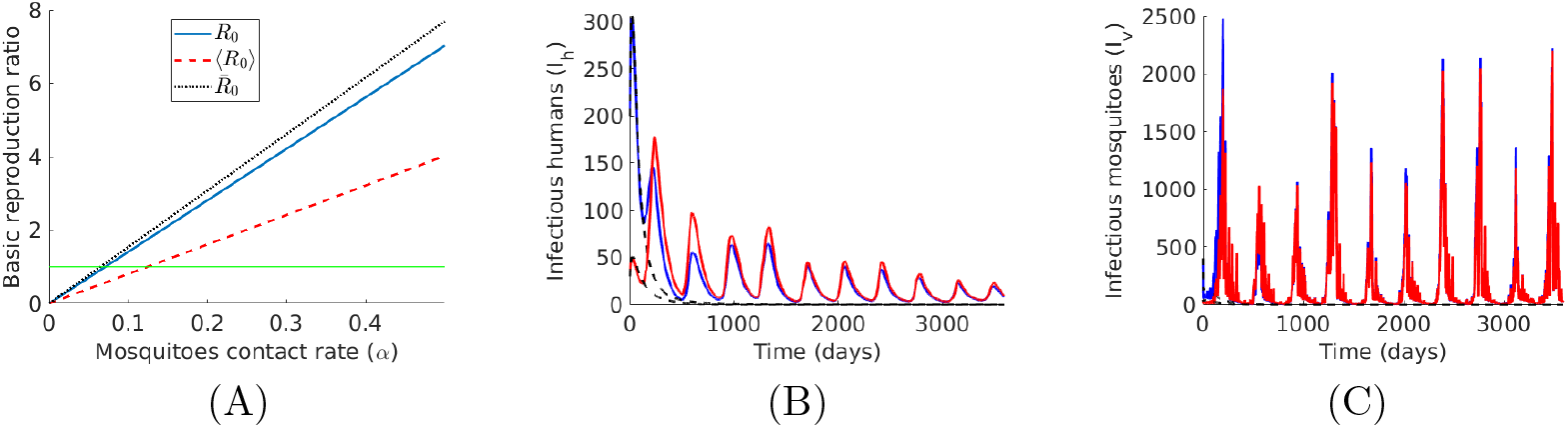
Here we set *ρ*_*h*_ = 0.01, *γ*_*h*_ = 0.01, *r* = 0.9 and Λ_*h*_ = 0.05. **A** Comparison between the basic reproduction number 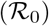, the average basic reproduction number 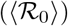 and the threshold parameter 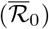, for range of the mosquitoes contact rate (*α*). **B-C** The long term dynamics of infectious human and infectious mosquitoes with *α* = 0.1. We use the temperature data of Nkomazi (South Africa) and find that 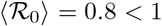; 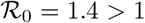. Other parameters are given by Tab. 3 and Tab. 2. We illustrate the behavior of the model using time dependent parameters *b*_*v*_, *μ*_*v*_, *β*_2_, *θ* (solid line) and average constant parameters ⟨*b*_*v*_⟩, ⟨*μ*_*v*_⟩, ⟨*β*_2_⟩, ⟨*θ*⟩ (dot line). The simulation is run with two initial data of model (2.1) given in Tab. 3.

### Illustration of disease extinction and persistence stated by Theorems 3.1 and 3.2

From Theorem 3.1, 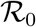 is a threshold parameter to determine whether malaria persists in the population. We choose the total number of human and mosquitoes at initial time (*t* = 0) to be *N*_*h*_(0) = 1000 and *N*_*v*_ (0) = 2000 respectively. Two initial data of the model are considered and given in Table 3. We take *ρ*_*h*_ = 0.01, *γ*_*h*_ = 0.01, *r* = 0.9, Λ_*h*_ = 0.05 and the other parameters are given by Table 2 and 3. By setting the mosquitoes contact rate per human host to *α* = 0.1 per day, numerical simulations complete the theoretical analysis that there exists a global attractive positive periodic solution when 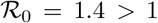 (Figure 2 B, C). With *α* = 0.06, Figure 3 supports the theoretical fact that the disease-free equilibrium *M*_0_ is globally asymptotically stable when 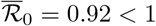. Note that, here we have 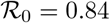.

**Figure 3:**
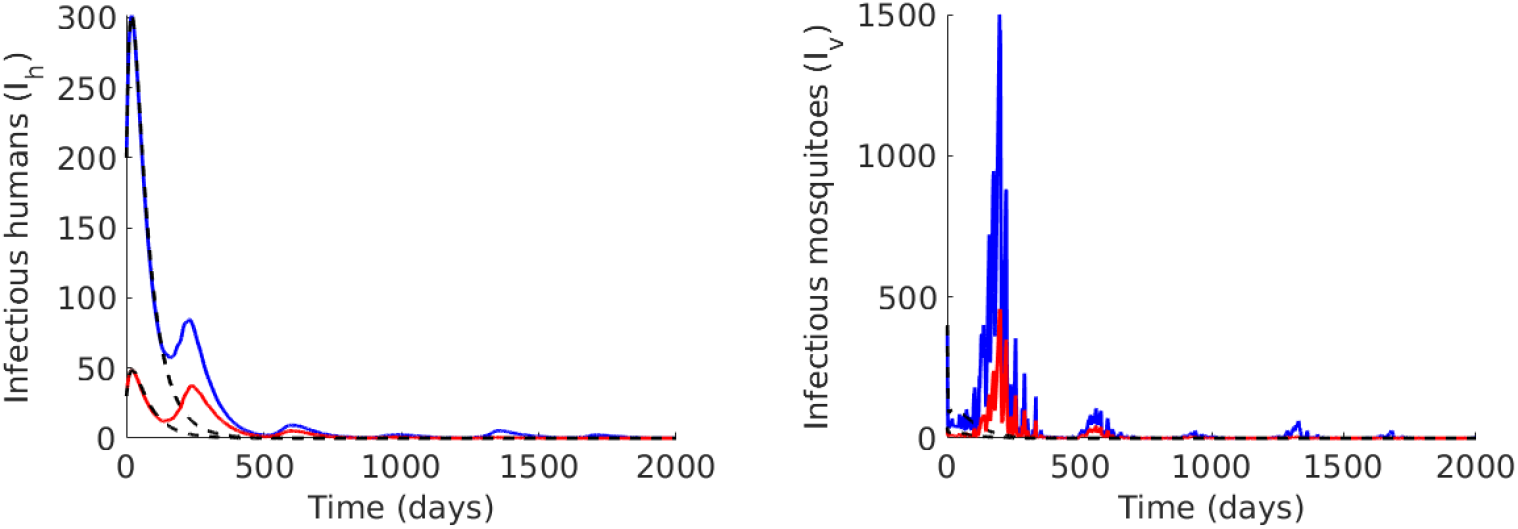
The long term behaviours (for two initial values) of four classes of population illustrated that the disease free state *M*_0_ is globally stable. Here, we use Λ_*h*_ = 0.05, *ρ*_*h*_ = 0.01, *γ*_*h*_ = 0.01, *r* = 0.9 and *α* = 0.06 per day and other parameters and initial values are given by Tab. 3 and Tab. 2. Then 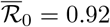, 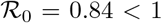 and 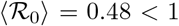. We illustrate the behavior of the model using time dependent parameters *b*_*v*_, *μ*_*v*_, *β*_2_, *θ* (solid line) and average constant parameters ⟨*b*_*v*_⟩, ⟨*μ*_*v*_⟩, ⟨*β*_2_⟩, ⟨*θ*⟩ (dotted line).

We now illustrate that model (2.1) can exhibits a backward bifurcation for some well chosen parameters, i.e., there can be stable endemic equilibrium when the basic reproduction number 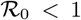. For a better illustration, here we choose the total number of human and mosquitoes at initial time (*t* = 0) to be *N*_*h*_(0) = 10 and *N*_*v*_ (0) = 20. Moreover, based on Figure 2A we choose *α* = 0.068 such that 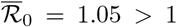 and 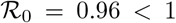. Then, Figure 4 illustrates the persistence of the epidemic although the 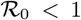.

**Figure 4:**
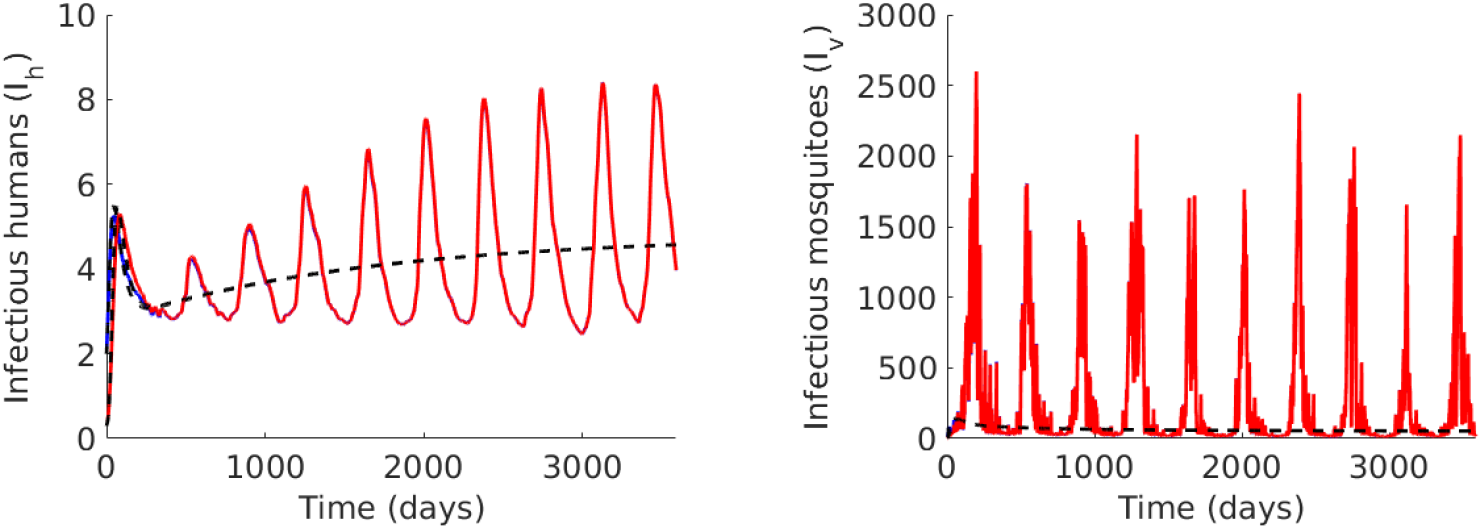
Illustration of the backward bifurcation of the model. Here, we use Λ_*h*_ = 0.05, *ρ*_*h*_ = 0.01, *γ*_*h*_ = 0.01, *r* = 0.9 and *α* = 0.068 per day such that 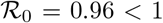 and 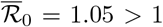. We also set *N*_*v*_ (0) = 20, *N*_*h*_(0) = 10 and other parameters are given by Tab. 3 and Tab. 2. We illustrate the behavior of the model using time dependent parameters *b*_*v*_, *μ*_*v*_, *β*_2_, *θ* (solid line) and average constant parameters ⟨*b*_*v*_⟩, ⟨*μ*_*v*_⟩, ⟨*β*_2_⟩, ⟨*θ*⟩ (dotted line).

### A case study

In this section, we estimate parameters of model (2.1)-(2.4) which are assumed to be variable (namely the human recruitment, disease induce mortality and recovery rate, Λ, *ρ*_*h*_, *γ*_*h*_ and *r*; the mosquitoes contact rate per human *α*) and the total number of human and mosquitoes at time *t* = 0, (*N*_*h*_(0), *N*_*v*_ (0)). We then study the transmission trend of malaria in Nkomazi, South Africa. Simulation results are given to show that our model with periodic parameters matches the seasonal fluctuation data reasonably well. The daily numbers of human malaria cases from the study region corresponds to the term *I*_*h*_(*t*) of model (2.1). Since we assume seven model parameters to be variable, *π* := (Λ, *ρ*_*h*_, *γ*_*h*_, *r*, *α*, *N*_*v*_(0), *N*_*h*_(0)), we then find the value 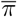 which minimize the difference (Δ[*π*]) between model prediction (*I*_*h*_) and the malaria cases of Nkomazi (*I*_cases_) from day *D*_*S*_ = October 1, 1997 to day *D*_*F*_ = December 31, 2005: 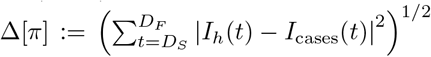. The value 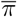 is identified with the MatLab nonlinear programming solver *fmincon*. Nkomazi malaria cases and the model fit well with 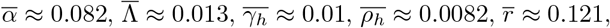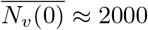 and 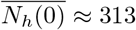 (see Figure 5).

**Figure 5:**
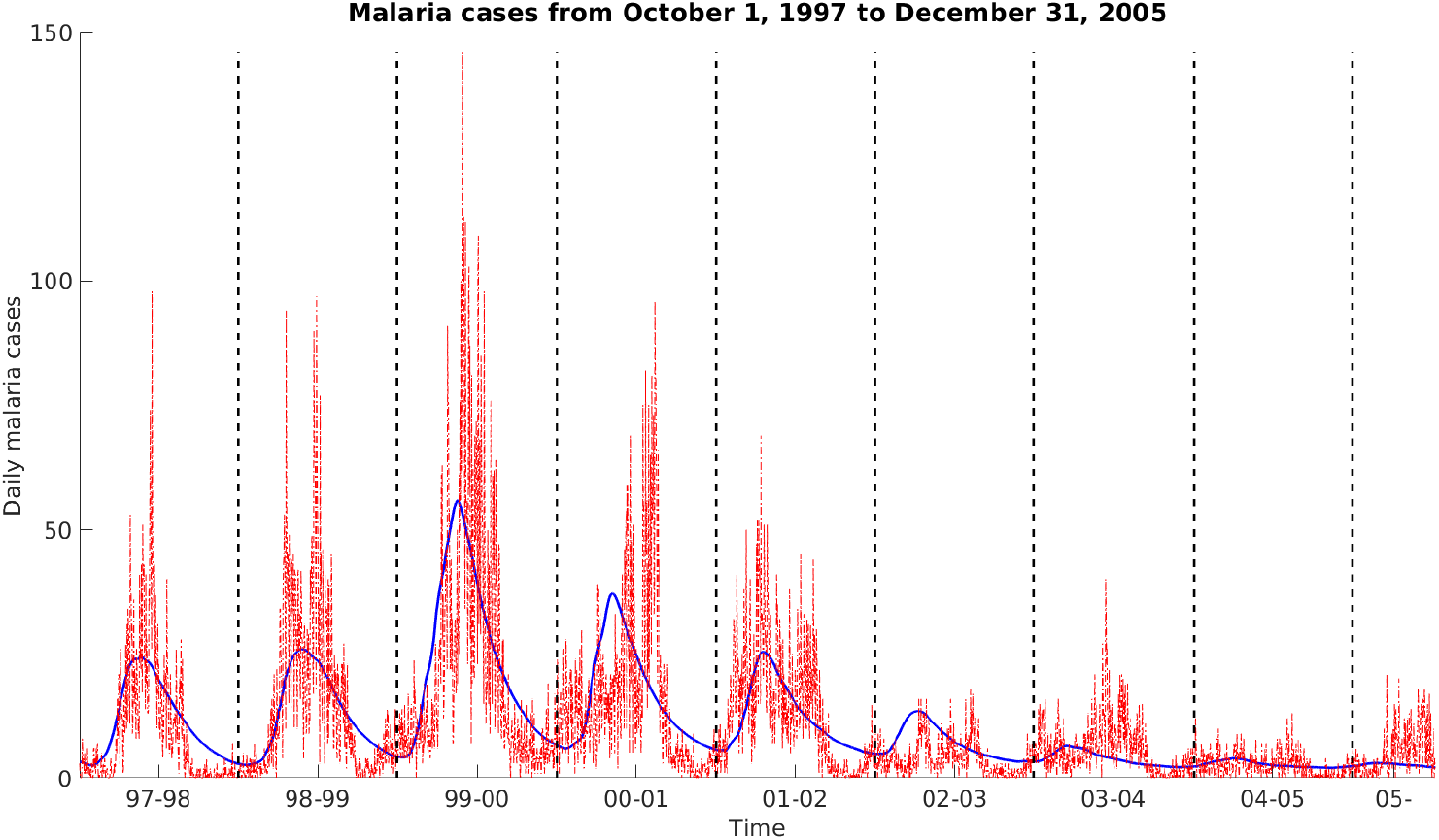
Daily malaria cases: reported number in Nkomazi (red dot line) and simulation curve (blue solid line) from October 1, 1997 to December 31, 2005. Here, we have a better fit with *α* ≈ 0.082, Λ ≈ 0.013, *γ*_*h*_ ≈ 0.01, *ρ*_*h*_ ≈ 0.0082, *r* ≈ 0.121, *N*_*v*_ (0) ≈ 2000 and *N*_*h*_(0) ≈ 313. We find 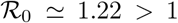 and 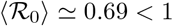. Other parameters are given by Tab. 3 and Tab. 2.

The simulation result based on our model exhibits the seasonal fluctuation and matches the data reasonably well. We estimate the basic reproduction number and the average basic reproduction number 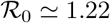 and 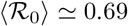 respectively. Furthermore, the value of 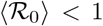 suggests that the epidemic is not endemic in Nkomazi leading to a wrong interpretation as illustrated by Figure 5.

## 5 Existence of the disease-free periodic state and preliminary results

The aim of this section is to derive some preliminary remarks on system (2.3)-(2.4). These results include the existence of the disease-free periodic state and the unique maximal non-autonomous semiflow associated with this system.

### 5.1 Existence of the disease-free *ω*-periodic state

The *N*_*h*_-equation of (3.5) gives

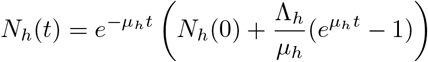

trough the arbitrary initial value *N*_*h*_(0), and has a unique periodic attractive solution in ℝ_+_, 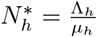. It follows that 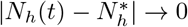 as *t* → ∞. Thus 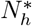 is globally attractive on ℝ_+_.

Concerning the *N*_*v*_-equation of (3.5), we have

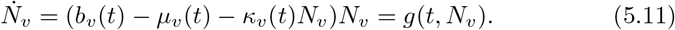

It is easy to see that 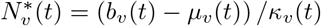 is a positive *ω*-periodic state of (5.11), ℝ_+_ is positively invariant for the above periodic equation and that and that *g* is strongly subhomogeneous in the sense that *g*(*t*, *νN*_*v*_) > *νg*(*t*, *N*_*v*_) for any *t* ≥ 0, *N*_*v*_ ∈ ℝ_+_ and *ν* ∈ (0, 1). By Assumption 2.1, item 2, equation (5.11) is such that 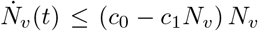, wherein *c*_*i*_ are positive constants. Thus, solutions of (5.11) are ultimately bounded in ℝ_+_. By Theorem 2.3.2 in [49] as applied to the Poincaré map associated with system (5.11), it follows that 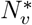 is the unique positive *ω*-periodic solution of (5.11), which is globally attractive in ℝ_+_.

### 5.2 Preliminary results

As in [25], the following vector order in ℝ^*n*^ will be used. For *u*, *v* ∈ ℝ^*n*^, we write

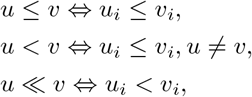

where *i* = 1, …, *n*.

#### Definition 5.1

*Consider two maps τ* : [0, ∞) × Ω → (0, ∞] *and* 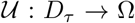, *where D*_*τ*_ = {(*t*, *s*, **v**) ∈ [0, ∞)^2^ × Ω : *s* ≤ *t* ≤ *s* + *τ* (*s*, **v**)}. *We say that* 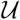 *is a* maximal non-autonomous semiflow on Ω *if U satisfies the following properties:*

i. 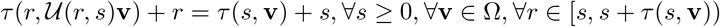.
ii. 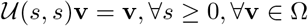.
iii. 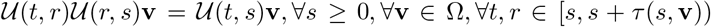 *with t* ≥ *r*.
iv. *If τ* (*s*, **v**) < +∞, *then* 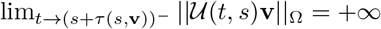.

We first derive that the Cauchy problem (2.3)-(2.4) generates a unique globally defined and positive non-autonomous semiflow.

#### Theorem 5.2

*Let Assumption 2.1 be satisfied. Then there exits a map τ* : [0, ∞)×Ω → (0, ∞] *and a maximal non-autonomous semiflow U* : *D*_*τ*_ → Ω, *such that for each x*_0_ := *x*(0) ∈ Ω *and each s* ≥ 0, 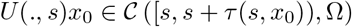 *is a unique maximal solution of* (2.3)-(2.4). *The map U* (*t, s*)*x*_0_ := (*x*_*h*_(*t*)^*T*^, *x*_*v*_(*t*)^*T*^)^*T*^ *satisfied the following properties: The subsets X*_0_ *and ∂X*_0_ *are both positively invariant under the non-autonomous semiflow U; in other words*,

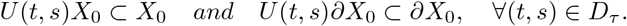

*Proof.* The proof of this result is rather standard. Standard methodology apply to provide the existence and uniqueness of the semiflow of system (2.3)-(2.4) [46, 24, 28, 34].

Let us check the positive invariance of Ω with respect to the semiflow *U* (*t*). To do so, let *x*_0_ = (*E*_*h*_(0), *I*_*h*_(0), *R*_*h*_(0), *E*_*v*_ (0), *I*_*v*_ (0), *N*_*h*_(0), *N*_*v*_ (0))^*T*^ ∈ Ω, we shall prove that *U* (*t, s*)*x*_0_ = (*E*_*h*_(*t*), *I*_*h*_(*t*), *R*_*h*_(*t*), *E*_*v*_ (*t*), *I*_*v*_ (*t*), *N*_*h*_(*t*), *N*_*v*_ (*t*))^*T*^ ∈ Ω for all (*t, s*) ∈ *D*_*τ*_. By Assumption 2.1 we find that *N*_*h*_(*t*) > 0 and *N*_*v*_ (*t*) > 0, for all *t* ≥ 0.

For the other variables, we consider first the case where *I*_*h*_(0) > 0. Using the continuity of the semiflow we find *t*_0_ > 0 such that *I*_*h*_(*t*) > 0 for *t* ∈ [0, *t*_0_]. If there is *t*_1_ ∈ [0, *t*_0_] such that *E*_*h*_(*t*_1_) + *I_h_*(*t*_1_) + *R_h_*(*t*_1_) = 1, then 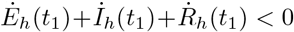. Therefore *E_h_*(*t*)+*I_h_*(*t*)+*R_h_*(*t*) ≤ 1 for all *t* ∈ [0, *t*_0_]. Similarly, *E*_*v*_ (*t*) + *I*_*v*_ (*t*) ≤ 1 for all *t* ∈ [0, *t*_0_]. If *E*_*v*_ (*t*_1_) = 0 for *t*_1_ ∈ [0, *t*_0_], then *E*_*v*_-equation of system (2.3) gives 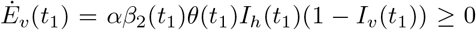. Thus *E*_*v*_ (*t*) ≥ 0 for all *t* ∈ [0, *t*_0_]. The same arguments give successively that *R_h_*(*t*) ≥ 0, *I*_*v*_ (*t*) ≥ 0 and *E*_*h*_(*t*) ≥ 0, for 0 < *t* ≤ *t*_0_.

We next show that *I_h_*(*t*) remains positive for all *t* ≥ *t*_0_. Proceeding by contradiction we suppose that *I_h_*(*t*) > 0 for 0 ≤ *t* < *t*_0_ and *I_h_*(*t*_0_) = 0. Then 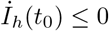. On the other hand, by *I_h_*-equation of System (2.3), 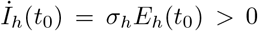, which is a contradiction. This complete the proof of the first part of theorem in the case *I_h_*(0) > 0. It remains to consider the case *I_h_*(0) = 0. In this case, recalling *E_h_*(0) + *R_h_*(0) ≤ 1 so that either *E_h_*(0) > 0 or *R_h_*(0) > 0. Without loss of generality, we suppose *E_h_*(0) > 0 and denote by *x*^*δ*^ (*t*) the solution (*δ*-solution) of System (2.3) with 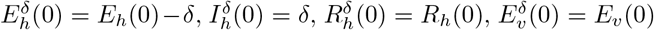, 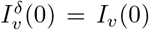, *N_h_*(0)^*δ*^ = *N_h_*(0), *N*_*v*_ (0)^*δ*^ = *N*_*v*_ (0); where 0 < *δ* < *E*_*h*_(0). By what we have already proved, the *δ*-solution *x*^*δ*^ (*t*) remains on Ω for all *t* > 0. Taking *δ* → 0, the first part of the theorem follows.

To end the proof of the theorem, let

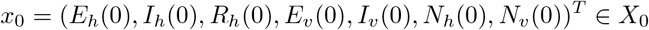

be given and let us denote for each (*t*, *s*) ∈ *D*_*τ*_,

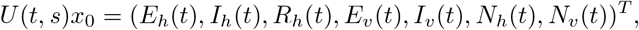

the orbit of system (2.3) passing through *x*_0_. Let us set *y_h_*(*t*) = *E_h_*(*t*) + *I_h_*(*t*) and *y*_*v*_ (*t*) = *E*_*v*_ (*t*) + *I*_*v*_ (*t*). It follows from system (2.3) that 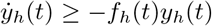 and 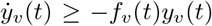; with *f_h_*(*t*) = *αβ*_1_(*t*)*θ*(*t*)*I*_*v*_*N*_*v*_/*N*_*h*_ + Λ_*h*_/*N*_*h*_ + *ρ*_*h*_ + *γ*_*h*_ and *f*_*v*_ (*t*) = *αβ*_2_(*t*)*θ*(*t*)*I*_*h*_ + *b*_*v*_ (*t*). That is

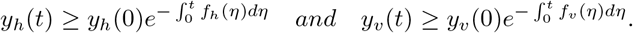

This end the proof of the fact that *U* (*t, s*)*X*_0_ ⊂ *X*_0_.

Now, let *x*_0_ ∈ *∂X*_0_. We have 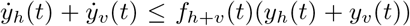, where *f*_*h*+*v*_ (*t*) = *αβ*_1_(*t*)*θ*(*t*)*N*_*v*_/*N*_*h*_ + *αβ*_2_(*t*)*θ*(*t*) + *ρ*_*h*_(*I*_*h*_ + *E_h_*). Then,

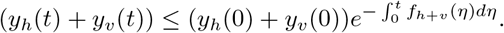

Since *y_h_*(0) + *y*_*v*_ (0) = 0, we find that *y_h_*(*t*) + *y*_*v*_ (*t*) = 0. Therefore, *U* (*t, s*)*∂X*_0_ ⊂ *∂X*_0_.

Recalling (3.6), we now deal with the spectral properties of the linearized system of model (2.3) at the disease-free equilibrium *M*_0_.

In the periodic case, we let *W*_λ_(*t*, *s*), *t* ≥ *s*, *s* ∈ ℝ, be the evolution operator of the linear *ω*-periodic system

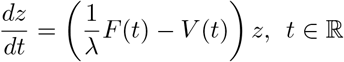

with parameter *λ* ∈ (0, ∞). Clearly, Φ_*F* − *V*_ (*t*) = *W*_1_(*t*, 0), ∀*t* ≥ 0. Note that 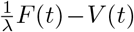 is cooperative. Thus, the Perron-Frobenius theorem (see [42], Theorem A.3) implies that *ρ*(*W_λ_*(*ω*, 0) is an eigenvalue of *W_λ_*(*ω*, 0) with nonnegative eigenvector. We can easily find that the matrix *W*_*λ*_(*s* + *ω, s*) is similar to the matrix *W_λ_*(*ω*, 0), and hence *σ*(*W_λ_*(*s* + *ω, s*)) = *σ*(*W_λ_*(*ω*, 0)) for any *s* ∈ ℝ, where *σ*(*D*) denotes the spectrum of the matrix *D*. It is easy to verify that system (2.3) satisfies assumptions (A1)-(A7) in [45]. Thus, recalling (3.8), we have the following results.

#### Lemma 5.3

*([45], Theorem 2.1). The following statements are valid:*

1. *If ρ*(*W_λ_*(*ω*, 0) = 1 *has a positive solution λ*_0_, *then λ*_0_ *is an eigenvalue of operator L, and hence* 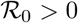.
2. *If* 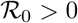, *then* 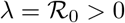 *is the unique solution of ρ*(*W_λ_*(*ω*, 0) = 1.
3. 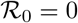 *if and only if ρ*(*W_λ_*(*ω*, 0) < 1 *for all λ* > 0.

Note that the above result can be used to numerically compute the reproduction number 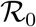 of the model, namely Lemma 5.3 item (ii).

The next result shows that 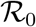 is a threshold parameter for the local stability of the disease-free periodic state *M*_0_ of system (2.3).

#### Lemma 5.4

*([45], Theorem 2.2). The following statements are valid:*

1. 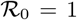 *if and only if ρ*(Φ_*F* −*V*_ (*ω*)) = 1.
2. 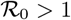 *if and only if ρ*(Φ_*F* −*V*_ (*ω*)) > 1.
3. 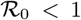 *if and only if ρ*(Φ_*F* −*V*_ (*ω*)) < 1.

Let *P* : Ω → Ω be the Poincaré map associated with system (2.3), that is 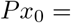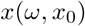 where *x*(*ω*, *x*_0_) is the unique solution of system (2.3) with *x*(0, *x*_0_) = *x*_0_. We easily find that *P*^*n*^*x*_0_ = *x*(*nω*, *x*_0_), for all *n* ≥ 0.

The following lemma will be useful later to derive the malaria persistence in the host population when the basic reproduction number 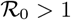.

#### Lemma 5.5

*If* 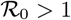, *then there exists ϵ* > 0, *such that for any* 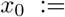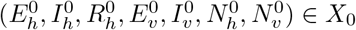 *with* ‖*x*_0_ − *M*_0_‖ ≤ *ϵ, we have*

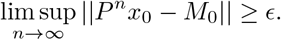

*Proof.* Let *η* > 0. By the continuity of the solutions with respect to the initial values, we find *ϵ* = *ϵ*(*η*) > 0 such that for all *x*_0_ ∈ *X*_0_ with ‖*x*_0_ − *M*_0_‖ ≤ *ϵ*, there holds ‖*x*(*t, x*_0_) − *x*(*t, M*_0_)‖ < *η*, ∀*t* ∈ [0, *ω*]. We further claim that

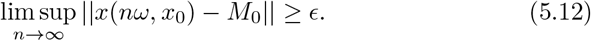

Assume by contradiction that (5.12) does not hold. Then without loss of generality, we assume that ‖*x*(*nω*, *x*_0_) − *M*_0_‖ < *ϵ*, for all *n* ≥ 0 and for some *x*_0_ ∈ *X*_0_. It follows that

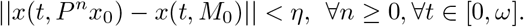

For any *t* ≥ 0, let *t* = *nω* + *s* and *n* is the largest integer less than or equal to *t/ω*. Therefore, we have

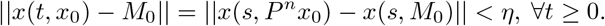

Recalling that *x*(*t*, *x*_0_) = (*E_h_*(*t*), *I_h_*(*t*), *R_h_*(*t*), *E*_*v*_ (*t*), *I*_*v*_ (*t*), *N_h_*(*t*), *N*_*v*_ (*t*))^*T*^, and since *x*(*t*, *M*_0_) ∈ *∂X*_0_ for all *t* ≥ 0 (see Theorem 5.2); it then follows that *E_h_*(*t*) < *η*, *I_h_*(*t*) < *η*, *E*_*v*_ (*t*) < *η*, *I*_*v*_ (*t*) < *η* for all *t* ≥ 0. By the *N_h_*-equation of (2.3), we have 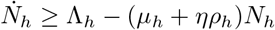. Note that the perturbed system

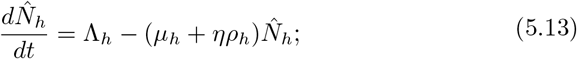

has a unique globally attractive (periodic) solution in ℝ_+_ defined by

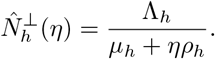

Since, 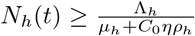 with *C*_0_ > 1 a constant, it comes 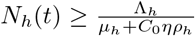, for *t* sufficiently large.

By the *R_h_*-equation of (2.3), we also have 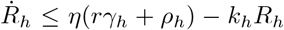, from where we find a constant *C*_1_ > 0 such that *R_h_*(*t*) ≤ *C*_1_*η*, for *t* sufficiently large. Further, by the globally attractivity in ℝ_+_ of 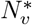 (see section 5.1), we also have 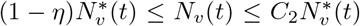, for sufficiently large *t* (with *C*_2_ > 1 a constant). Therefore, the term 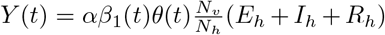 is such that 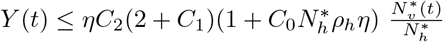, for sufficiently large *t*.

Now, let us find an estimation of the term *N*_*v*_ (*t*)*/N*_*h*_(*t*). Again, we have 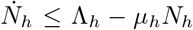. From where, 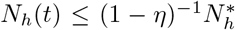, for *t* sufficiently large, and so 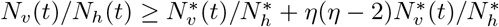.

Then, from the *E_h_*, *I_h_*, *E_v_* and *I_v_* equations of system (2.3), we obtain, for sufficiently large time *t*,

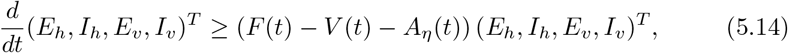

with

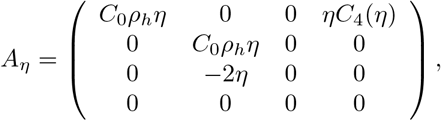

with 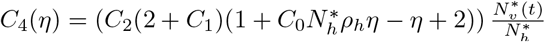.

Since 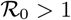, Lemma 5.4 implies that *ρ*(Φ_*F* −*V*_ (*ω*) > 1. We can choose *η* > 0 small enough such that 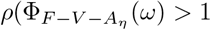.

We then consider the following system

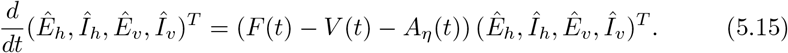

By [48], Lemma 2.1, it follows that there exists a positive *ω*-periodic function 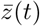 such that 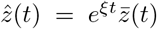 is a solution of system (5.15), with 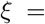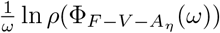. Since 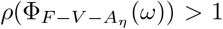, *ξ* is a positive constant. Let *t* = *nω* and *n* be nonnegative integer and get

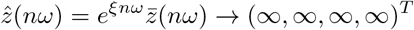

as *n* → ∞, since *ωξ* > 0 and 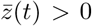. For any nonnegative initial condition (*E_h_*(0), *I_h_*(0), *E*_*v*_ (0), *I*_*v*_ (0)) of system (5.14), there exists a sufficiently small *n*_0_ > 0 such that 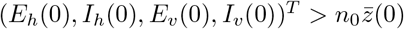. By the comparison principle (Theorem B.1, [42]) we have 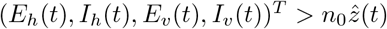, for all *t* > 0. Thus, we obtain (*E_h_*(*nω*), *I_h_*(*nω*), *E*_*v*_ (*nω*), *I*_*v*_ (*nω*))^*T*^ → (∞, ∞, ∞, ∞)^*T*^ as *n* → ∞, a contradiction with the first part of Theorem 5.2.

## 6 Proof of Theorems 3.1 and 3.2

This section is devoted to the threshold dynamics results state by Theorems 3.1 and 3.2.

### 6.1 Proof of Theorem 3.1

From Lemma 5.4, it follows that the disease-free equilibrium for System (2.3)-(2.4) is locally asymptotically stable if 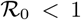 and unstable if 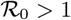, which ends the proof of Theorem 3.1 (i). In the sequel, we check the disease persistence results. It remains to check item (ii) of the theorem.

By Theorem 5.2, the discrete system *P* = {*P*^*n*^}_*n*∈ℕ_ admits a global attractor in Ω. For any 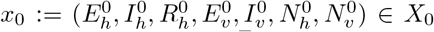, Let *x*(*t*, *x*_0_) = (*E_h_*(*t*), *I_h_*(*t*), *R_h_*(*t*), *E*_*v*_ (*t*), *I*_*v*_ (*t*), *N*_*h*_(*t*), *N*_*v*_ (*t*))^*T*^ be the orbit of (2.3) passing through *x*_0_. We have show that, both Ω, *X*_0_ and *∂X*_0_ are positively invariant with respect to the non-autonomous semiflow *U* (Theorem 5.2). Clearly, *∂X*_0_ is relatively closed in Ω, and there is exactly one fixed point 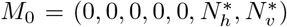 of *P* in *∂X*_0_.

Note that the non-linear system

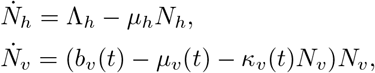

admits a global asymptotic equilibrium 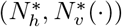, see section 5.1. Then, Lemma 5.5 implies that 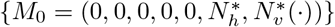 is an isolated invariant set in Ω and *W^s^*({*M*_0_}) ∩ *X*_0_ = ∅. We can also note that every orbit in *∂X*_0_ approches to *M*_0_ and *M*_0_ is acyclic in *∂X*_0_. By [49], Theorem 1.3.1, it follows that *P* is uniformly persistence with respect to the pair (*X*_0_, *∂X*_0_). That is, there exists a *δ* > 0 such that any solution *x*(*t*, *x*_0_) of system (2.3) with initial value *x*_0_ ∈ *X*_0_ satisfies lim inf_*t*→∞_ *d*(*x*(*t*, *x*_0_), *∂X*_0_) ≥ *δ*.

Furthermore, [49], Theorem 1.3.6, implies that the discrete system {*P^n^*}_*n*∈ℕ_ has a fixed point 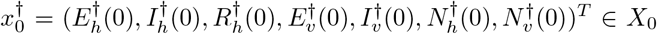. Then, by (*E_h_*, *I*_*h*_, *R*_*h*_, *E*_*v*_, *I*_*v*_)-equation of system (2.3) and the irreducibility of the cooperative matrix 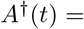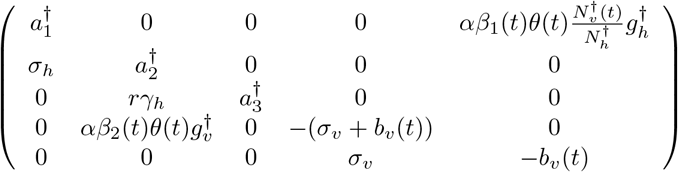, it follows that 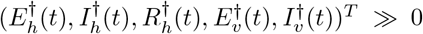 for all *t* ≥ 0. Where 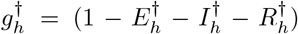 and 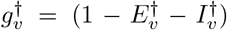; wherein 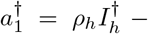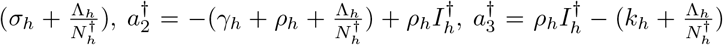. Therefore, 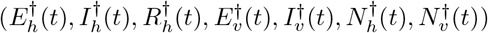 is a positive *ω*-periodic solution of system (2.3). This end the proof of the second part of Theorem 3.1.

### 6.2 Proof of Theorem 3.2

By the *N_h_*-equation of system (2.3), it comes 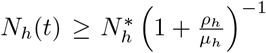 for all *t* > 0 sufficiently large. Further, due to the globally attractivity in ℝ_+_ of 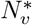 (see section 5.1), we have 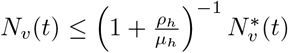 for all *t* sufficiently large. Therefore,

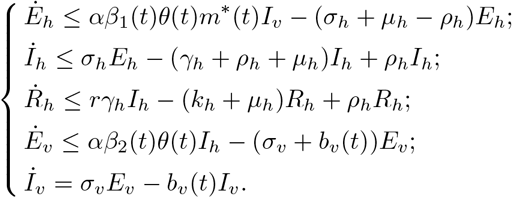

So, for all *t* sufficiently large, we obtain

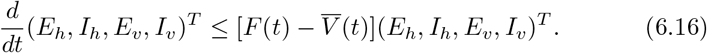

Now, we introduce the solution *z* of

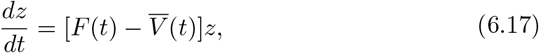

where *z*(0) is given.

Applying Lemma 5.4, we know that 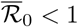 if and only if 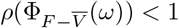. By [48], Lemma2.1, it follows that there exists a positive *ω*-periodic function 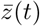 such that 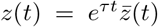 is a solution of system (6.17), with 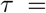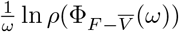. Since 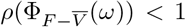, then *τ* is a negative constant; and we have *z*(*t*) → 0 as *t* → +∞. This implies that the zero solution of system (6.17) is globally asymptotically stable. For any non-negative initial value (*E_h_*(0), *I_h_*(0), *E*_*v*_ (0), *I*_*v*_ (0)) of system (6.16), there is a sufficiently large *C*_0_ > 0 such that (*E_h_*(0), *I_h_*(0), *E*_*v*_ (0), *I*_*v*_ (0)) ≤ *C*_0_*z*(0). Then, the comparison principle (Theorem B.1, [42]) gives (*E_h_*(0), *I_h_*(0), *E*_*v*_ (0), *I*_*v*_ (0)) ≤ *C*_0_*z*(*t*), for all *t* > 0, where *C*_0_*z*(*t*) is also the solution of system (6.16). We then get (*E_h_*(*t*), *I_h_*(*t*), *E*_*v*_ (*t*), *I*_*v*_ (*t*))^*T*^ → (0, 0, 0, 0)^*T*^ as *t* → +∞. It is easy to find that 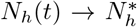 as *t* → +∞. Let *ε* > 0, we find *t_ε_* > 0 such that *I_h_*(*t*) ≤ *ε* for all *t* ≥ *t_ε_*. Then, the *R_h_*-equation of (2.3) gives 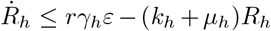, for large time *t*. From where *R_h_*(*t*) → 0 as *t* → +∞. This ends the proof of Theorem 3.2.

### 6.3 We always have 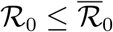

Let *Y*_*V*_ (*t, s*), *t* ≥ *s*, the evolution operator of the linear *ω*-periodic system (3.7) associated to the matrix *V* (*t*), with *Y*_*V*_ (*s, s*) = *I* (the identity matrix). We find that,

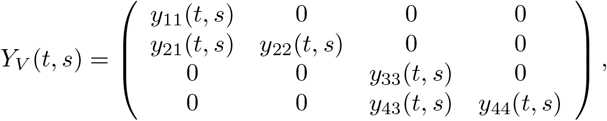

with 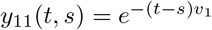; 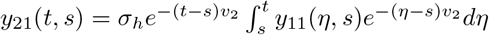; 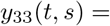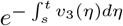; 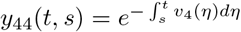; 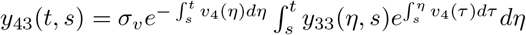; and where *v*_1_ = *σ*_*h*_ + *μ*_*h*_; *v*_2_ = *γ*_*h*_ + *ρ*_*h*_ + *μ*_*h*_; *v*_3_(*t*) = *σ*_*v*_ + *b*_*v*_ (*t*); *v*_4_(*t*) = *b*_*v*_ (*t*).

By the same way we, the evolution operator 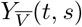 of the linear *ω*-periodic system (3.7) associated to the matrix 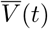, with 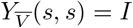 is given by

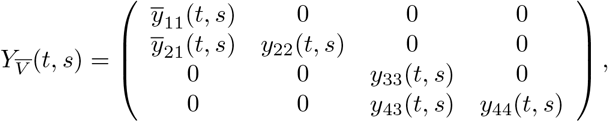

with 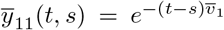; 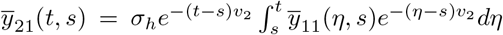 and 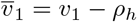.

Further, we can find *J* > 0 and *l* > 0 such that ‖*Y*_*V*_ (*t, s*)‖ ≤ *Je*^−*l*(*t*−*s*)^ and 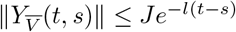 for all *t* ≥ *s*. Therefore,

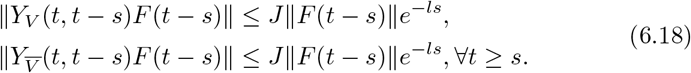

Recall operators *L*_*V*_ and 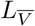 defined by (3.8) with the matrix *V* and 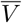 respectively. Clearly, *L*_*V*_ and 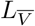 are positive operators. From (6.18) we easily get that *L*_*V*_ and 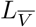 are bounded, and so, continuous on 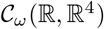. Since 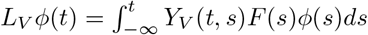 it comes

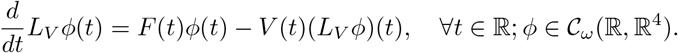

Then for any positive constant *c* > 0, there exists *U* = *U* (*c*) > 0 such that 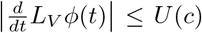 for all *t* ∈ [0, *ω*], 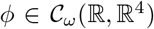 with ‖*ϕ*‖ ≤ *c*. Hence, the Ascoli–Arzela theorem gives the compacity of *L*_*V*_ on 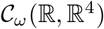. By the same arguments, we obtain the compacity of 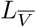.

Recall that spectral radius of operators *L*_*V*_ and 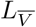 are 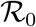 and 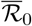 respectively. Obviously, when *ρ*_*h*_ = 0, we have 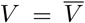, *i.e.* 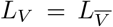 and then 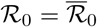. Now let *ρ*_*h*_ > 0. Then 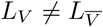 and we have 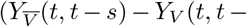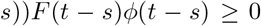 for all 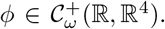. From the primitivity (or non-supporting) property of operators *L_V_* and 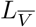 its comes that 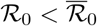 when *ρ_h_* > 0. See [32, 40] for some development on positive operator theory.

## Acknowledgements

GJA acknowledges the support of the Fogarty International Center (FIC) of the National Institutes of Health (NIH) under Award Number D43TW009343 and the University of California Global Health Institute (UCGHI).

## References

[1] Abdelrazec, A., Gumel, A. B. (2017). Mathematical assessment of the role of temperature and rainfall on mosquito population dynamics. Journal of mathematical biology, 74(6), 1351–1395.

[2] Abiodun, G.J., Okosun, O.K., Witbooi, P.J., Maharaj, R (2018). Exploring the impact of climate variability on malaria transmission using a dynamic mosquito-human malaria model. The Open Infectious Diseases Journal. https://www.benthamopen.com/EPUB/BMS-TOIDJ-2018-17.

[3] Abiodun, G. J., Witbooi, P. J., Okosun, K. O. (2017). Mathematical modelling and analysis of mosquito-human malaria model. International Journal of Ecological Economics and Statistics™, 38(3), 1–22.

[4] Abiodun, G. J., Maharaj, R., Witbooi, P. & Okosun, K. O. (2016). Modelling the influence of temperature and rainfall on the population dynamics of Anopheles arabiensis. Malaria journal, 15(1), 364.

[5] Adeola, A. M., Botai, O. J., Olwoch, J. M., Rautenbach, C. D. W., Adisa, O. M., Taiwo, O. J., Kalumba, A. M. (2016). Environmental factors and population at risk of malaria in N komazi municipality, South Africa. Tropical Medicine and International Health, 21(5), 675–686.

[6] Agusto, F.B., Gumel, A.B. and Parham, P.E., 2015. Qualitative assessment of the role of temperature variations on malaria transmission dynamics. Journal of Biological Systems, 23(04), p.1550030.

[7] Alonso, D., Bouma, M. J., Pascual, M. (2011). Epidemic malaria and warmer temperatures in recent decades in an East African highland. Proceedings of the Royal Society of London B: Biological Sciences, 278(1712), 1661–1669.

[8] Anderson, R. M., May, R. M. (1992). Infectious diseases of humans: dynamics and control. Oxford university press.

[9] Aron, J. L., May, R. M. (1982). The population dynamics of malaria. In The population dynamics of infectious diseases: theory and applications (pp. 139–179). Springer, Boston, MA.

[10] Bacaër, N. (2007). Approximation of the basic reproduction number *R*_0_for vector-borne diseases with a periodic vector population. Bulletin of mathematical biology, 69(3), 1067–1091.

[11] Bakary, T., Boureima, S., Sado, T. (2018). A mathematical model of malaria transmission in a periodic environment. Journal of biological dynamics, 12(1), 400–432.

[12] Beck-Johnson, L. M., Nelson, W. A., Paaijmans, K. P., Read, A. F., Thomas, M. B., Bjørnstad, O. N. (2013). The effect of temperature on Anopheles mosquito population dynamics and the potential for malaria transmission. PLOS one, 8(11), e79276.

[13] Beier, J. C. (1998). Malaria parasite development in mosquitoes. Annual review of entomology, 43(1), 519–543.

[14] Blayneh, K., Cao, Y., Kwon, H. D. (2009). Optimal control of vector-borne diseases: treatment and prevention. Discrete and Continuous Dynamical Systems B, 11(3), 587–611.

[15] Bray, R. S., Garnham, P. C. C. (1982). The life-cycle of primate malaria parasites. British Medical Bulletin, 38(2), 117–122.

[16] Briere, J. F., Pracros, P., Le Roux, A. Y., Pierre, J. S. (1999). A novel rate model of temperature-dependent development for arthropods. Environmental Entomology, 28(1), 22–29.

[17] Burkot, T. R., Graves, P. M., Paru, R., Battistutta, D., Barnes, A., Saul, A. (1990). Variations in malaria transmission rates are not related to anopheline survivorship per feeding cycle. The American journal of tropical medicine and hygiene, 43(4), 321–327.

[18] Chitnis, N., Cushing, J. M., Hyman, J. M. (2006). Bifurcation analysis of a mathematical model for malaria transmission. SIAM Journal on Applied Mathematics, 67(1), 24–45.

[19] Chitnis, N., Hyman, J. M., Cushing, J. M. (2008). Determining important parameters in the spread of malaria through the sensitivity analysis of a mathematical model. Bulletin of mathematical biology, 70(5), 1272.

[20] Craig, M. H., Snow, R. W., le Sueur, D. (1999). A climate-based distribution model of malaria transmission in sub-Saharan Africa. Parasitology today, 15(3), 105–111.

[21] Dembele, B., Friedman, A., Yakubu, A. A. (2009). Malaria model with periodic mosquito birth and death rates. Journal of biological dynamics, 3(4), 430–445.

[22] Eikenberry, S. E., Gumel, A. B. (2018). Mathematical modeling of climate change and malaria transmission dynamics: a historical review. Journal of mathematical biology, 1–77.

[23] Friedman, A. (2013). Epidemiological models with seasonality. In Mathematical Methods and Models in Biomedicine (pp. 389–410). Springer, New York, NY.

[24] Hartman P, Ordinary Differential Equations. John Wiley and Sons, New-York, 1964.

[25] Hirsch, M. W., Smith, H. (2006). Monotone dynamical systems. In Handbook of differential equations: ordinary differential equations (Vol. 2, pp. 239–357). North-Holland.

[26] Hoshen, M. B., Morse, A. P. (2004). A weather-driven model of malaria transmission. Malaria Journal, 3(1), 32.

[27] Ikeda, T., Behera, S.K., Morioka, Y., Minakawa, N., Hashizume, M., Tsuzuki, A., Maharaj, R. and Kruger, P., 2017. Seasonally lagged effects of climatic factors on malaria incidence in South Africa. Scientific reports, 7(1), p.2458.

[28] Katok, A., Hasselblatt, B. (1997). Introduction to the modern theory of dynamical systems (Vol. 54). Cambridge university press.

[29] Lafferty, K. D. (2009). The ecology of climate change and infectious diseases. Ecology, 90(4), 888–900.

[30] Laneri, K., Paul, R. E., Tall, A., Faye, J., Diene-Sarr, F., Sokhna, C., … RodÓo, X. (2015). Dynamical malaria models reveal how immunity buffers effect of climate variability. Proceedings of the National Academy of Sciences, 112(28), 8786–8791.

[31] Liu, L., Zhao, X. Q., Zhou, Y. (2010). A tuberculosis model with seasonality. Bulletin of Mathematical Biology, 72(4), 931–952.

[32] Marek, I. (1970). Frobenius theory of positive operators: Comparison theorems and applications. SIAM Journal on Applied Mathematics, 19(3), 607–628.

[33] Mordecai, E. A., Paaijmans, K. P., Johnson, L. R., Balzer, C., Ben-Horin, T., de Moor, E., … Lafferty, K. D. (2013). Optimal temperature for malaria transmission is dramatically lower than previously predicted. Ecology letters, 16(1), 22–30.

[34] Morris, W. Hirsch, Smale, S. (1973). Differential equations, dynamical systems and linear algebra. Academic Press college division.

[35] Ngwa, G. A., Shu, W. S. (2000). A mathematical model for endemic malaria with variable human and mosquito populations. Mathematical and Computer Modelling, 32(7-8), 747–763.

[36] Nwankwo, A., & Okuonghae, D. (2019). Mathematical assessment of the impact of different microclimate conditions on malaria transmission dynamics. Mathematical Biosciences and Engineering, 16(3): 1414–1444.

[37] Okuneye, K., Gumel, A. B. (2017). Analysis of a temperature-and rainfall-dependent model for malaria transmission dynamics. Mathematical biosciences, 287, 72–92.

[38] Paaijmans, K. P., Read, A. F., Thomas, M. B. (2009). Understanding the link between malaria risk and climate. Proceedings of the National Academy of Sciences, 106(33), 13844–13849.

[39] Roy, M., Bouma, M., Dhiman, R. C., Pascual, M. (2015). Predictability of epidemic malaria under non-stationary conditions with process-based models combining epidemiological updates and climate variability. Malaria journal, 14(1), 419.

[40] Sawashima, I.(1964), On spectral properties of some positive operators, Nat. Sci. Report Ochanomizu Univ., 15, 53–64.

[41] Silal, S. P., Barnes, K. I., Kok, G., Mabuza, A., Little, F. (2013). Exploring the Seasonality of Reported Treated Malaria Cases in Mpumalanga, South Africa. PLoS ONE, 8(10), e76640.

[42] Smith, H. L., Waltman, P. (1995). The theory of the chemostat: dynamics of microbial competition (Vol. 13). Cambridge university press.

[43] Tumwiine, J., Mugisha, J.Y.T. and Luboobi, L.S., 2007. A mathematical model for the dynamics of malaria in a human host and mosquito vector with temporary immunity. Applied Mathematics and Computation, 189(2), pp.1953–1965.

[44] Van den Driessche, P., Watmough, J. (2008). Further notes on the basic reproduction number. In Mathematical epidemiology (pp. 159–178). Springer, Berlin, Heidelberg.

[45] Wang, W., Zhao, X. Q. (2008). Threshold dynamics for compartmental epidemic models in periodic environments. Journal of Dynamics and Differential Equations, 20(3), 699–717.

[46] Wanner, G., Hairer, E. (1991). Solving ordinary differential equations II. Stiff and Differential-Algebraic Problems.

[47] World Health Organization (2017). World Malaria Report 2017. http://www.who.int/malaria/publications/world-malaria-report-2017/en/ (Accessed: June 2018)

[48] Zhang, F., Zhao, X. Q. (2007). A periodic epidemic model in a patchy environment. Journal of Mathematical Analysis and Applications, 325(1), 496–516.

[49] Zhao, X.Q. (2003) Dynamical Systems in Population Biology. Springer-Verlag, New York.

